# Combining tRNA sequencing methods to characterize plant tRNA expression and post-transcriptional modification

**DOI:** 10.1101/790451

**Authors:** Jessica M. Warren, Thalia Salinas-Giegé, Guillaume Hummel, Nicole L. Coots, Joshua M. Svendsen, Kristen C. Brown, Laurence Maréchal-Drouard, Daniel B. Sloan

## Abstract

Differences in tRNA expression have been implicated in a remarkable number of biological processes. There is growing evidence that tRNA genes can play dramatically different roles depending on both expression and post-transcriptional modification, yet sequencing tRNAs to measure abundance and detect modifications remains challenging. Their secondary structure and extensive post-transcriptional modifications interfere with RNA-seq library preparation methods and have limited the utility of high-throughput sequencing technologies. Here, we combine two modifications to standard RNA-seq methods by treating with the demethylating enzyme AlkB and ligating with tRNA-specific adapters in order to sequence tRNAs from four species of flowering plants, a group that has been shown to have some of the most extensive rates of post-transcriptional tRNA modifications. This protocol has the advantage of detecting full-length tRNAs and sequence variants that can be used to infer many post-transcriptional modifications. We used the resulting data to produce a modification index of almost all unique reference tRNAs in *Arabidopsis thaliana*, which exhibited many anciently conserved similarities with humans but also positions that appear to be “hot spots” for modifications in angiosperm tRNAs. We also found evidence based on northern blot analysis and droplet digital PCR that, even after demethylation treatment, tRNA-seq can produce highly biased estimates of absolute expression levels most likely due to biased reverse transcription. Nevertheless, the generation of full-length tRNA sequences with modification data is still promising for assessing differences in relative tRNA expression across treatments, tissues or subcellular fractions and help elucidate the functional roles of tRNA modifications.

## INTRODUCTION

Despite their central function in cell physiology and the increasing interest in quantifying their expression, transfer RNAs (tRNAs) remain difficult to sequence using conventional RNA sequencing (RNA-seq) methods. tRNAs are poorly suited to high-throughput sequencing library preparation protocols for two reasons. First, tRNAs are extensively post-transcriptionally modified, and certain base modifications occurring at the Watson-Crick face stall or terminate reverse transcription (RT) through interference with base pairing or steric hindrance, effectively blocking cDNA synthesis (Motorin et al. 2007). The majority of RNA-seq protocols require that RNA is reverse transcribed into cDNA before sequencing, but RT of tRNAs often results in truncated cDNA products that lack 5’ adapters and are never sequenced. Second, RT for RNA-seq is normally primed off of a ligated 3’ adapter, but mature tRNAs have compact secondary and tertiary structure with tightly base-paired 5’ and 3’ termini, which can prevent adapter ligation (Wilusz 2015). Hybridization methods such as microarrays and northern blot analysis have been used to quantify multiple tRNA species, but these approaches require prior knowledge of an organism’s tRNA repertoire, and they may not able to evaluate expression levels of near-identical tRNAs because of cross-hybridization (Shigematsu et al. 2017). Additionally, RNA-seq has higher sensitivity for genes that are very highly or lowly expressed and can detect a much larger range of expression (Wang et al. 2009).

One significant technological advance in developing effective tRNA-seq protocols was the discovery that certain tRNA base modifications could be removed prior to RT using the demethylating enzyme AlkB from *Escherichia coli*. AlkB was shown to remove 1-methyladenine and 3-methylcytosine damage in single- and double-stranded DNA (Trewick et al. 2002) and later utilized by Zheng et al. (2015) and Cozen et al. (2015) to remove some RT-inhibiting modifications present on tRNAs. In these studies, tRNA-seq libraries treated with AlkB had sustainably higher abundance and diversity of tRNA reads. This was followed by the development of mutant forms of AlkB engineered to target specific modifications that appeared to be recalcitrant to demethylation by wild type AlkB treatment (Zheng et al. 2015; Dai et al. 2017). Although it is still not possible to remove all RT-inhibiting modifications with AlkB, the use of this enzyme represented a large step forward in tRNA sequencing.

In order to overcome inefficient adapter ligation, multiple methods have been developed involving modified adapter ligations (Pang et al. 2014; Smith et al. 2015; Zhong et al. 2015; Shigematsu et al. 2017) or a template-switching thermostable group II intron reverse transcriptase that eliminates the adapter ligation step entirely [DM-TGIRT] (Zheng et al. 2015). In the case of YAMAT-seq (Shigematsu et al. 2017), Y-shaped DNA/RNA hybrid adapters are utilized to specifically bind to the unpaired discriminator base and the -CCA sequence motif added to the 3’ end of all mature tRNAs. The utility of adapter protocols that utilize CCA-complementarity and ligation of both the 5’ and 3’ adapters to intact RNAs is that full-length, mature tRNAs are preferentially sequenced. In contrast, methods that ligate adapters in RNA/DNA step-wise fashion (Pang et al. 2014) or eliminate adapter ligation in the case of DM-TGIRT have been effective at broadly detecting tRNA gene transcription, but largely capture tRNA fragments. These truncated sequence reads are then difficult to confidently map to a single gene, requiring additional predictive models and bioinformatic solutions (Gogakos et al. 2017; Hoffmann et al. 2018; Torres et al. 2019). Despite the progress being made with AlkB to remove inhibitory modifications and methods such as YAMAT-seq to specifically target full-length tRNAs, no study has yet applied both approaches to combine the benefits of each.

Although post-transcriptional tRNA modifications have hindered efforts to sequence tRNAs, they are a universal and fundamental aspect of tRNA biology (Rojas-Benitez et al. 2015; Lyons et al. 2018). Base modifications are involved in tRNA folding and stability (Sampson and Uhlenbeck 1988; Vermeulen et al. 2005; Phizicky and Alfonzo 2010), translational accuracy and reading-frame maintenance (Urbonavicius et al. 2001; Hou et al. 2015), and tRNA fragment generation (Lyons et al. 2018; Huber et al. 2019). Not surprisingly given these fundamental roles, they have been implicated in numerous diseases (Phizicky and Hopper 2015), and there has been a long historical interest to identify, map, and quantify these modifications (Kuchino et al. 1987). Traditional methods to identify base modifications involved tRNA purification and either direct sequencing and fingerprinting or complete digestion of RNAs into nucleosides followed by liquid chromatography-mass spectrometry (LC-MS) (Kowalak et al. 1993; Ross et al. 2016). Soon after the discovery of reverse transcriptase enzymes, signatures of RT inhibition were also recognized as a means to map base modification positions (Youvan and Hearst 1979). Typical protocols involved primer extension assays and ^32^P-labeled DNA primers to detect blocked or paused primer extension signals (Schwartz and Motorin 2017). RT-based RNA modification detection assays were later extended through the use of chemical reagents that reacted with specific modified nucleotides to further confirm the identity of the modifications (Motorin et al. 2007). The advent of high-throughput sequencing has now facilitated the generation of entire maps or “indexes” of the modifications that cause RT-misincorporations across all tRNAs found in a species (Clark et al. 2016). These signatures include fragment generation because of RT termination but also read-through misincorporations, i.e. base misincorporation and indels produced from the RT enzyme reading through a modified base (Kietrys et al. 2017; Motorin and Helm 2019). Despite these advances, modification indexes have been produced for few species, and there has been limited investigation of tRNA modification profiles of isodecoders (tRNAs that share the same anticodon but have sequence variation at other parts along the tRNA). In addition, the function of some modifications is still unknown, and new modifications are still being discovered using a combination of RT-based modification mapping and mass spectrometry (Kimura et al. 2019).

tRNA gene organization, expression, and modification patterns affect a wide diversity of biological processes, and the role of individual tRNAs has seen increasing interest in the past twenty years (Hummel et al. 2019). Species commonly have conserved multigene isoacceptor families (a group of tRNAs that are acylated with the same amino acid but can have different anticodons) (Bermudez-Santana et al. 2010), and there is growing evidence that certain tRNA genes have functionally unique roles (Kondo et al. 1990; Goodarzi et al. 2016). One evolutionary lineage that has particularly complex tRNA metabolism is angiosperms, or flowering plants. Plant nuclear tRNAs can be present in numerous copies, some of which are organized into large, tandemly repeated arrays (Theologis et al. 2000; Cognat et al. 2013). Furthermore, the generation and function of tRNA-derived fragments (tRFs) has come under increasing scrutiny because their presence has been directly linked to plant cell growth and response to stresses (Cognat et al. 2017; Park and Kim 2018; Soprano et al. 2018). Plants also have a high degree of tRNA compartmentalization and trafficking because of the presence of two endosymbiotically derived organelles, the plastid and the mitochondria. Each organelle has its own set of tRNAs, but plant mitochondria are exceptional compared to bilaterian animals in that they require extensive import of nuclear-encoded tRNAs into the mitochondrial matrix to maintain translation (Salinas-Giege et al. 2015). Additionally, plants have been shown to have some of the most post-transcriptionally modified tRNAs identified to date (Iida et al. 2009; Machnicka et al. 2014). These modifications are increasingly being assigned new biological roles, making plants an ideal group to study the function of tRNA modifications.

Unfortunately, the reasons that make plant tRNA biology so fascinating (i.e. complex metabolism and extreme modification rates) have historically hindered investigation because of difficulties in applying high throughput sequencing and mapping methods, and recent advances in tRNA-seq such as the use of AlkB and CCA-complementary adapters have not yet been applied to plant systems. Here, we have combined these two tRNA-seq methods to detect full-length tRNAs, map them to unique reference gene sequences and identify RT-induced sequence variants in four different angiosperm species, focusing on the model system *Arabidopsis thaliana*.

## RESULTS

### YAMAT-seq captures mature, full-length tRNAs from the majority of *A. thaliana* tRNA reference genes

tRNAs were sequenced with a combination of two complementary methods, YAMAT-seq (Shigematsu et al. 2017) and AlkB treatment (Cozen et al. 2015; Zheng et al. 2015) to generate full-length, mature tRNA reads from *A. thaliana* leaf total-cellular RNA. The demethylating enzyme AlkB was used in conjunction with YAMAT-seq’s Y-shaped adapters that are complementary to the CCA motif found at the 3’ end of mature tRNAs to generate Illumina libraries composed almost entirely of tRNA sequences, with representatives from most genes in the reference *A. thaliana* database (all sequences in the database can be found in **supp. Table 1** with the corresponding tRNA gene identifiers). Greater than 99% of the reads mapped to a tRNA sequence when BLASTed to an *A. thaliana* tRNA database (e-value cutoff of 1e-6) (**supp. Table 2**). The remaining reads were largely derived from cytosolic/plastid rRNAs and adapter artifacts (**supp. Table 3**). Over 98% of the reads had a 3’-terminal CCA sequence. Although the proportion of reads that matched the reference sequence length varied among isoacceptor families (**Figure 1**), the vast majority of the reads were 74-95 nt after adapter trimming (representing the expected range of mature tRNA sequence lengths in *A. thaliana*), indicating that our library size selection successfully targeted full-length tRNAs. A small fraction of the reads, even those with a CCA tail, were truncated. These fragments (defined as having less than 70% hit coverage to a reference tRNA) appear to be largely generated from a subset of tRNAs, particularly certain nuclear tRNA-Thr and tRNA-Ala genes. Many of these fragments appeared to involve sequences that inverted and re-primed off of short regions of sequence similarity on the complementary strand, producing a “U-turn” molecule.

**Fig. 1.**
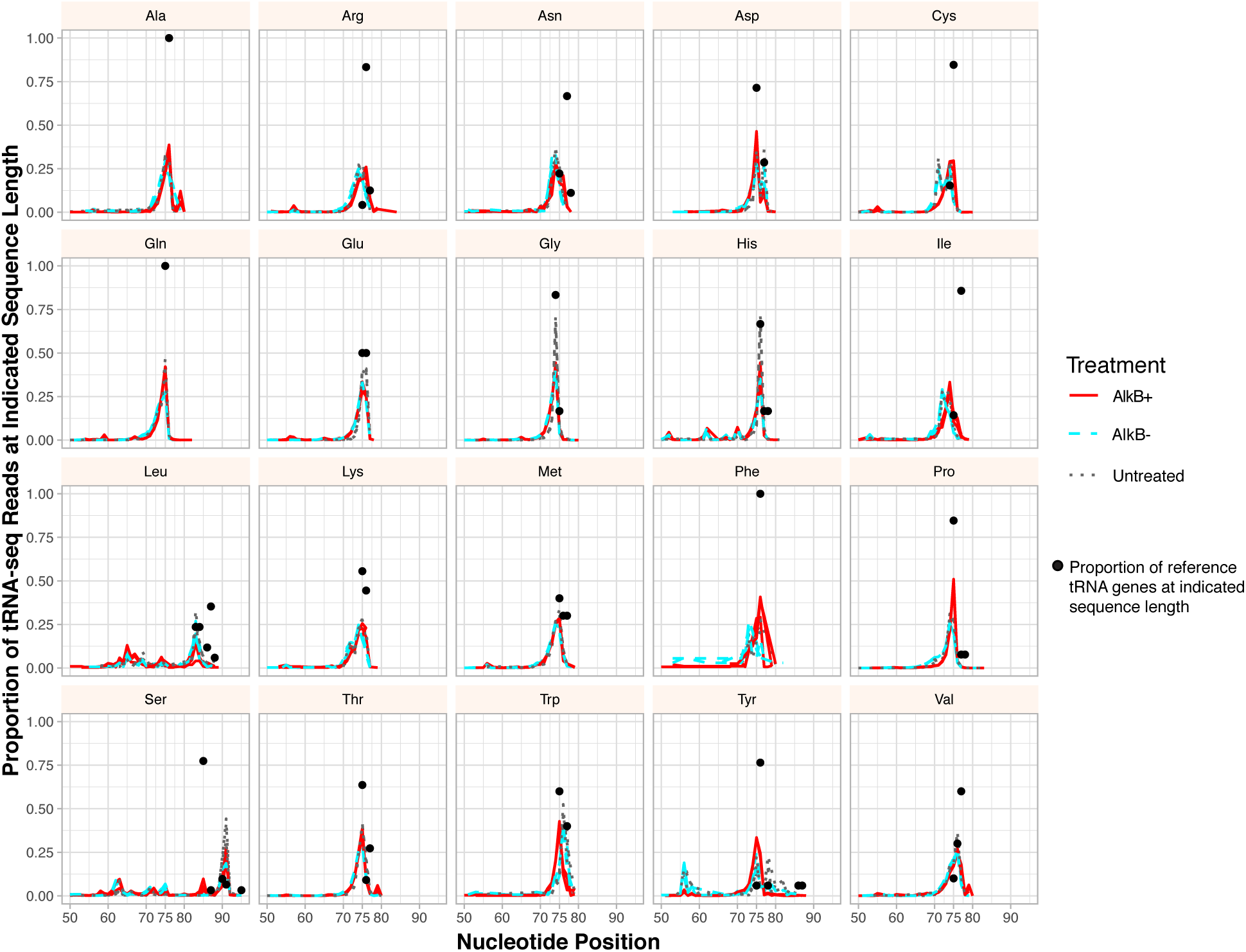
The proportion of tRNA-seq reads at different lengths based on isoacceptor family. Black dots represent the proportion of isoacceptor reference sequences at indicated length (e.g. 100% of the Ala-tRNA reference genes are 76 nt in length). Three biological replicates are plotted for each treatment (AlkB+, AlkB- and untreated). Note that the dominant peak for read length often corresponds to the length of the reference sequence(s).

### AlkB treatment improves detection of *A. thaliana* reference tRNAs

Base modifications known to stall or terminate RT are prevalent in tRNAs (Motorin et al. 2007). The demethylating enzyme AlkB has been shown to effectively remove some of these modifications (N^1^-methyladenosine [m^1^A], N^3^-methylcytosine [m^3^C]), thereby making certain tRNAs more amenable to sequencing (Cozen et al., 2015, Zheng et al., 2015). Our preliminary observations from testing AlkB treatments found a considerable effect on *A. thaliana* RNA integrity, with the majority of degradation occurring with exposure to the AlkB reaction buffer, regardless of whether the AlkB enzyme was included. Therefore, we used two types of negative controls in our experimental design to isolate the effects of AlkB from the buffer alone. Total RNA was either treated with the AlkB enzyme in the reaction buffer (AlkB+), treated with the reaction buffer alone (AlkB-), or left entirely untreated prior to RT. Libraries sequenced after the reaction buffer treatment alone and those that were entirely untreated did not substantially differ from each other in reference sequence detection and frequency (**supp. Figure 1**). However, treatment with AlkB resulted in better representation of the majority of cytosolic tRNAs and a moderate increase in the detection of some organellar tRNAs (**Figure 2, supp. Table 4**). Only 95-115 of the 183 nuclear tRNA reference genes were detected in untreated and AlkB-libraries (**supp. Table 5**), whereas AlkB+ treatment increased this range to 138-149. These reference coverage counts do not include tRNA reference genes that were exclusively detected with reads that were an equally good match to another reference sequence, which was the case for 4-13 genes per library (**supp. Table 5**). All 30 plastid and 17 of the 19 mitochondrial reference tRNAs were detected in at least one library with one mitochondrial tRNA-SerTGA gene only detected in a single AlkB+ library. No reads were detected for the mitochondrial genes tRNA-SerGCT-3360 and tRNA-SerGGA-3359. Ser and Tyr isoacceptors were the most likely to be undetected or have very low abundance in all libraries. Interestingly, three mitochondrial tRNA genes that are homologous to tRNA-PheGAA but have a mutated GTA (Tyr) anticodon and were previously annotated to be pseudogenes in the *A. thaliana* mitochondrial genome (GenBank: NC_037304.1) were detected as mature, expressed tRNAs in this analysis, which is consistent with earlier detection of expressed copies of one of these genes(Chen et al. 1997).

**Fig. 2.**
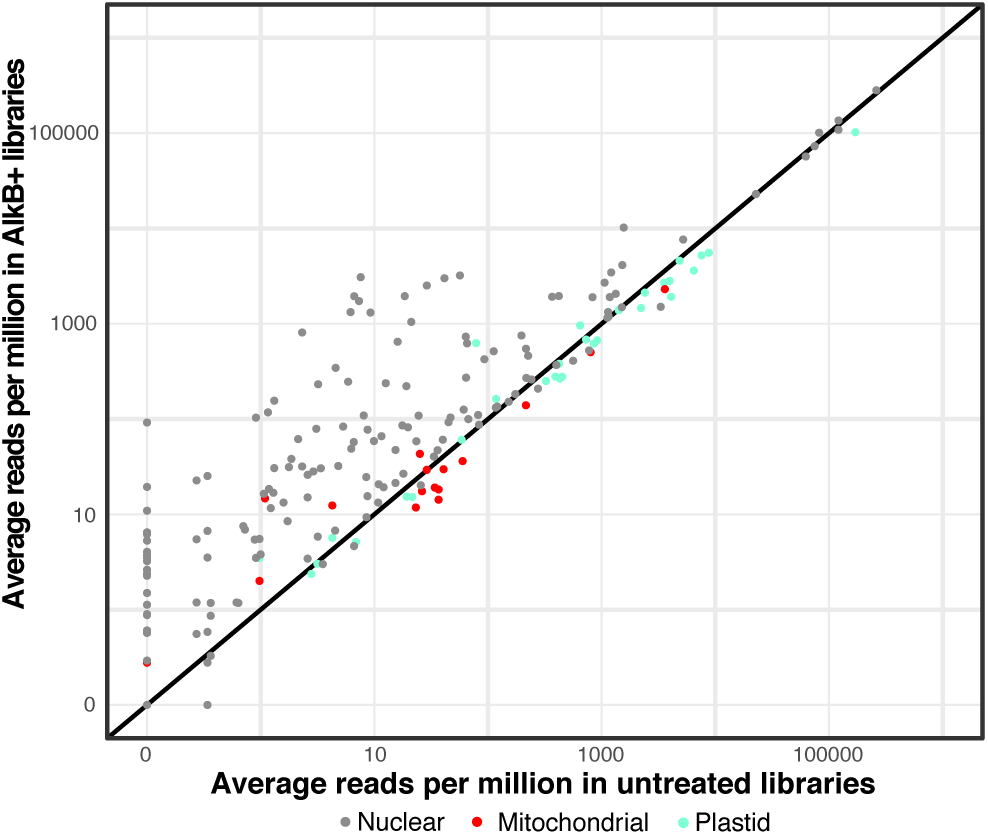
Demethylation treatment with AlkB increases the proportion of tRNA-seq reads for many tRNAs (points falling above the 1:1 line). Average read counts per million across three biological replicates are shown for each unique *A. thaliana* tRNA reference sequence in AlkB+ vs. untreated libraries.

In addition to the wild type AlkB, an AlkB mutant (D135S) has been specifically engineered to remove the modification N^1^-methylguanosine (m^1^G), which is known to inhibit RT activity (Zheng, et al. 2015). We performed tRNA-seq on three additional *A. thaliana* total-cellular RNA samples treated with either wild type AlkB, or a 2:1 ratio of wild type AlkB and D135S AlkB to test for further improvements in tRNA detection. We found only a moderate increase in the detection of a few genes when performing differential expression analysis on libraries treated with D135S (**supp. Table 6**). One D135S library did have a single read for a tRNA-TyrGTA gene that was undetected in the wild type libraries, but otherwise there was no increase in the number of genes detected in D135S libraries (**supp. Figure 1).**

### tRNA-seq profiles are dominated by nuclear tRNA-Pro and plastid tRNA-GlyGCC genes in four angiosperm species

The number of reads mapped to each reference tRNA sequence varied drastically and was heavily skewed towards multiple nuclear tRNA-Pro genes and a plastid tRNA-GlyGCC **(Figure 3)**. Together, nuclear tRNAs-Pro and the plastid tRNA-GlyGCC sequences comprised 86-90% of all reads in AlkB+ libraries and 91-93% in untreated and AlkB-libraries. To test whether this dominance of tRNA-Pro and tRNA-GlyGCC reads was unique to *A. thaliana* or a more widespread pattern in flowering plants, tRNAs were sequenced from leaf total-cellular RNA from another rosid (*Medicago truncatula*), an asterid (*Solanum tuberosum*) and a monocot (*Oryza sativa*), using the same tRNA-seq method described above with wild type AlkB. The resulting reads from all three species showed a similarly extreme skew towards nuclear tRNA-Pro and plastid tRNA-GlyGCC (**Figure 4)**.

**Fig. 3.**
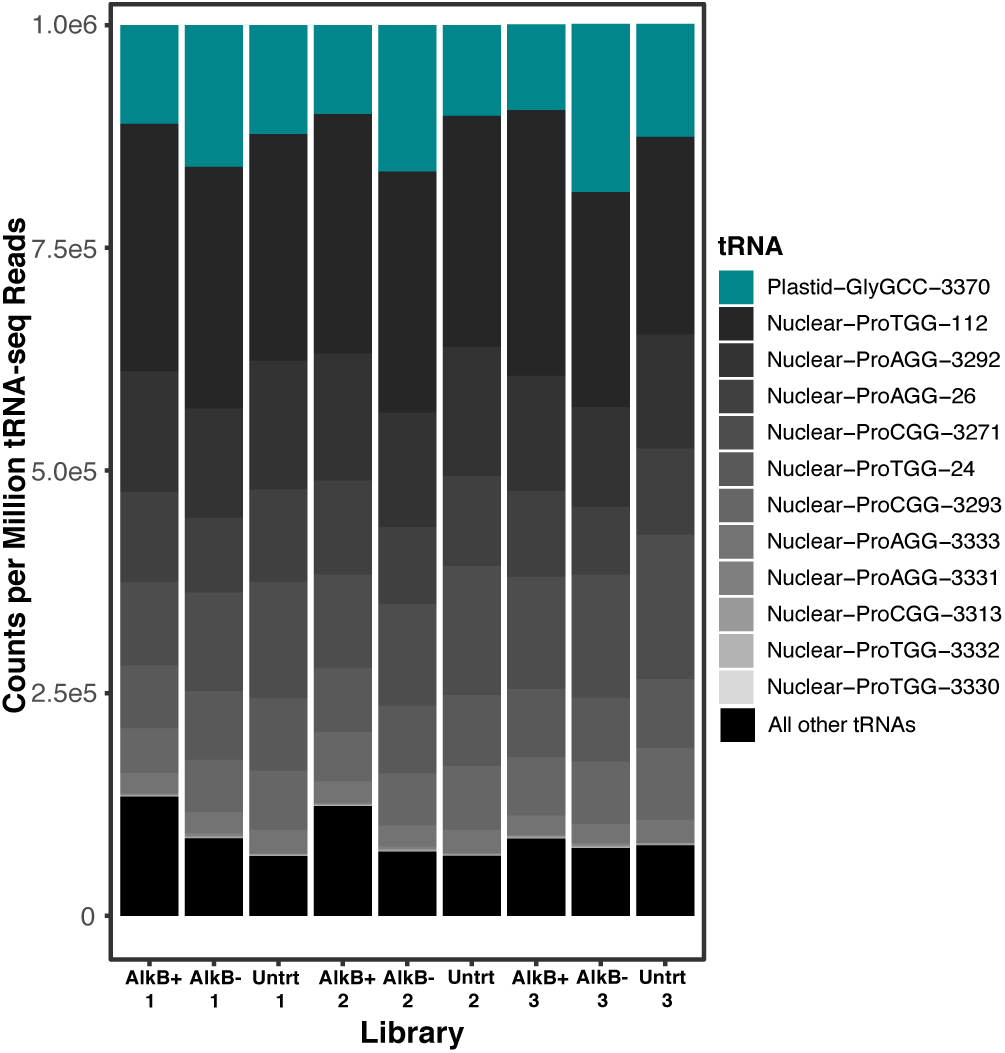
tRNA-seq read abundance in *A. thaliana* tRNA is dominated by nuclear tRNA-Pro and plastid tRNA-GlyGCC genes. All nuclear tRNA-Pro genes are shown and indicated by gray colors, with the darkest grays indicating the highest abundance, and the plastid tRNA-GlyGCC is indicated by teal. All other tRNAs have been grouped and shown in black.

**Fig. 4.**
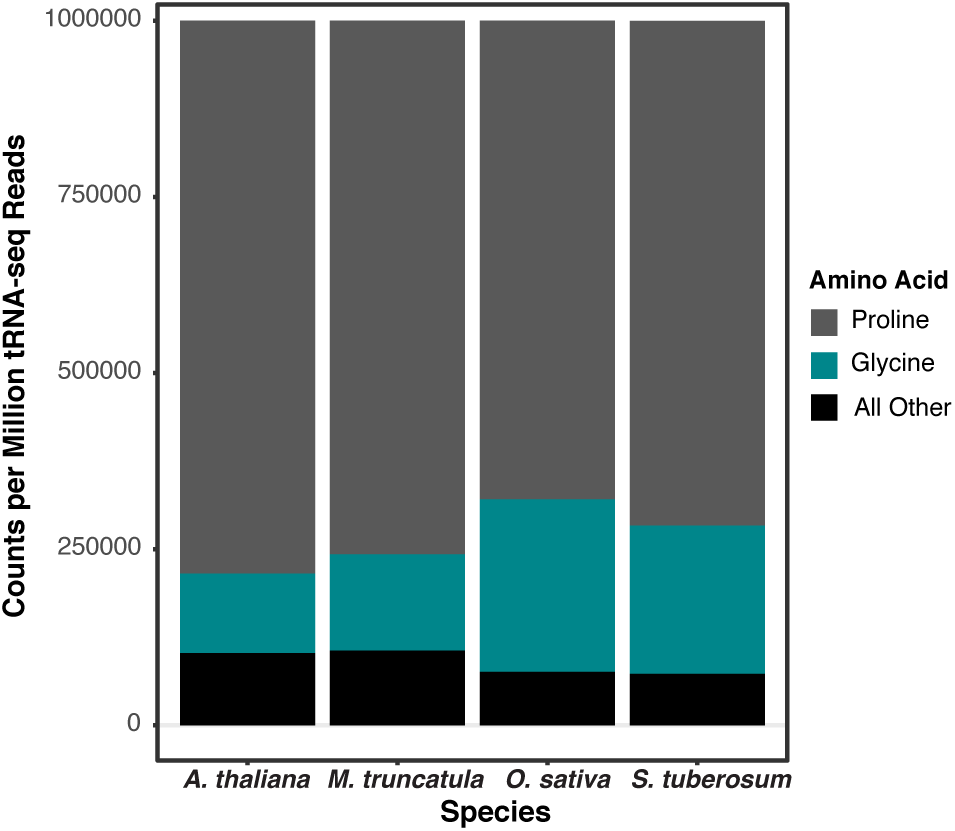
tRNA-seq read abundance is similarly dominated by only two isoacceptor families (tRNA-Pro and tRNA-Gly) in four angiosperm species.

### Persistent reverse transcription bias contributes to tRNA-seq coverage variation

The dominance of multiple nuclear tRNA-Pro genes and a plastid tRNA-GlyGCC gene was surprising because tRNA expression is generally expected to reflect codon usage after accounting for base-pairing modifications (Novoa et al. 2012). One possible artifactual source of variation in tRNA-seq read abundance is biased adapter ligation during library construction (Fuchs et al. 2015). In order to determine the abundance of tRNA-derived cDNA molecules independent of adapter ligation, droplet digital PCR (ddPCR) was performed on reverse transcribed, unligated subsamples of the three original *A. thaliana* RNA replicates using internal primers for four tRNA genes (**supp. Table 7**). There was a strong correlation between counts per million tRNA-seq reads (CPM) and ddPCR copies per nanogram (*p* = 0.03 and adjusted R^2^ = 0.91; **Figure 5**). These data suggest that adapter ligation bias is not the primary determinant of tRNA-seq coverage skew as cDNA copy number in ddPCR is reflective of the number of final tRNA-seq Illumina reads.

**Fig. 5.**
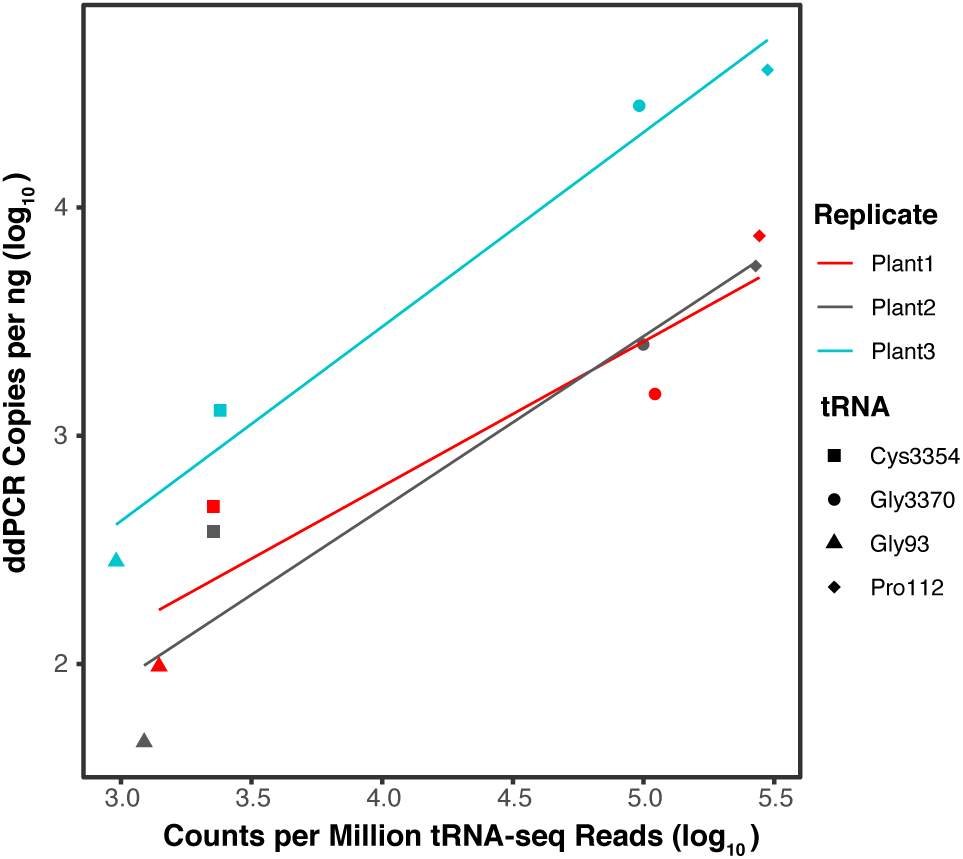
Droplet digital PCR (ddPCR) copies correlate with number of tRNA-seq reads. ddPCR copies per ng of cDNA plotted against counts per million tRNA-seq reads for four *A. thaliana* tRNA genes. Adjusted R^2^ values for separate linear regressions on biological replicates 1, 2, and 3 were 0.79, 0.85, and 0.96, respectively. When data points were averaged across biological replicates, linear regression yielded an adjusted R^2^ of 0.91 and a *p*-value of 0.03.

An additional source of sequencing bias could result from preferential RT of tRNAs with fewer modifications or less inhibitory secondary structures. In order to quantify tRNA abundance in total-cellular RNA without the intermediate step of RT, northern blot analysis was performed by probing for four *A. thaliana* tRNAs representing a range of CPM values from the tRNA-seq data. All four labeled probes were hybridized to three RNA replicates as well as a dilution series of a complementary oligonucleotide in order to quantify the tRNA signal (**supp. Table 8**). The tRNA-Pro gene with the highest CPM showed the weakest hybridization signal, and the highly expressed plastid tRNA-GlyGCC did not have the strongest intensity of the two plastid tRNAs probed (**Figure 6)**. The concentration estimates from the northern blot analysis present a truly striking contrast with the tRNA-seq data because the read abundance for the sampled tRNA-ProTGG was approximately 70,000-fold higher than one of the other sampled nuclear tRNAs (tRNA-SerCGA). This massive incongruence strongly points to biased tRNA RT (even after AlkB treatment) as contributing to the extreme coverage variation found in our tRNA-seq data.

**Fig. 6.**
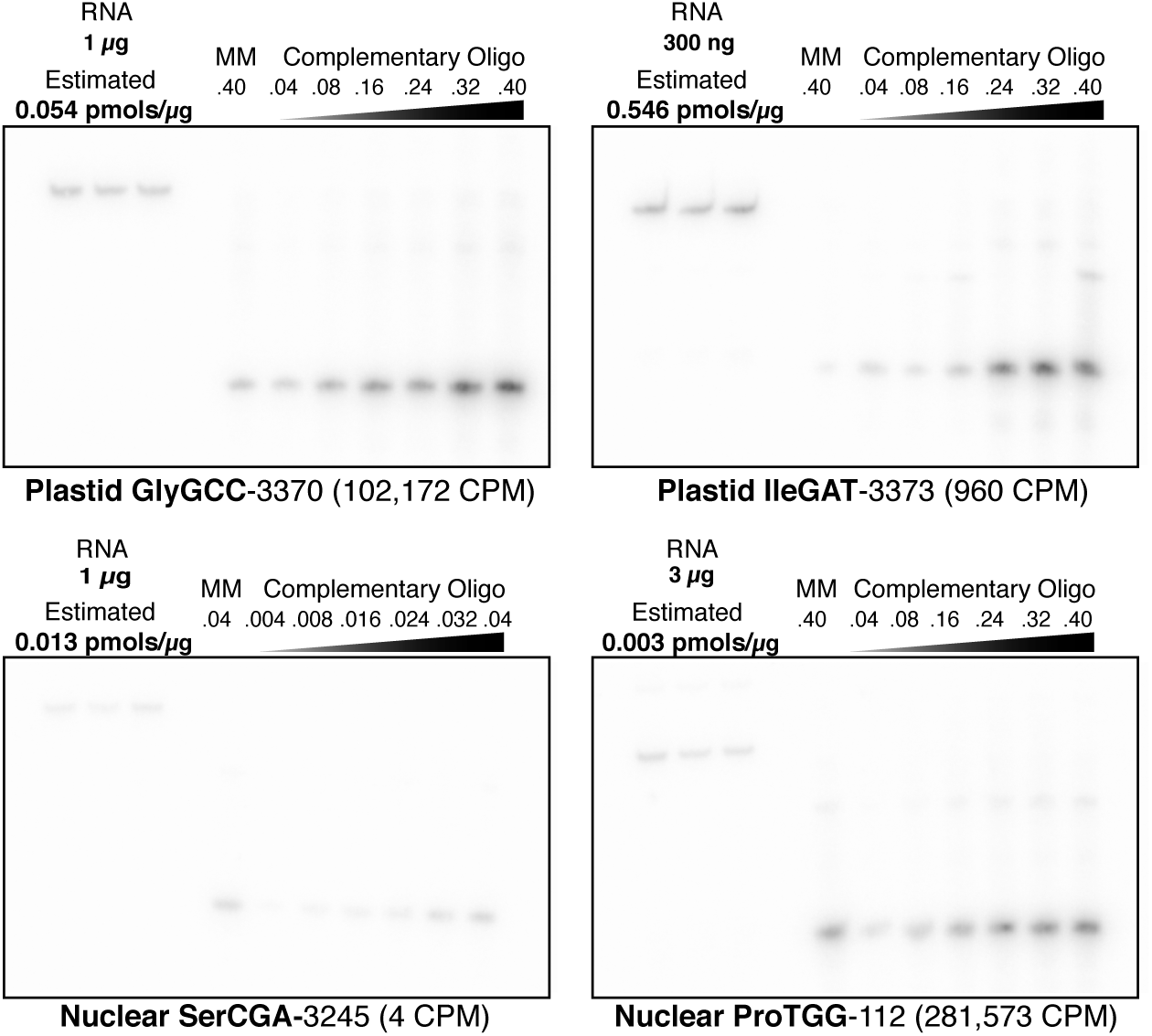
Northern blot analysis does not show the same expression dominance of nuclear tRNA-Pro and plastid tRNA-GlyGCC that was observed with tRNA-seq. Four different *A. thaliana* tRNA genes were probed from total cellular RNA and quantified. Each tRNA target membrane had three replicates of total *A. thaliana* RNA, which was quantified using a dilution series of a synthesized 38-nt oligonucleotide complementary to the corresponding probe. A single lane of a 38-nt mismatched oligonucleotide (MM), which had one or two non-complementary nucleotides relative to the probe, was also included on each membrane to test for cross hybridization of probes. The amounts of total RNA and oligonucleotides loaded on each blot were varied according to expected hybridization signal based on preliminary analyses. The mass of RNA and pmols of oligonucleotides are indicated above each corresponding lane. The estimated concentration of the target tRNA based on analysis of signal intensity in ImageJ is reported as pmols per μg of input RNA. The average Illumina tRNA-seq read counts per million (CPM) is indicated parenthetically for each tRNA.

Because tRNAs are multicopy genes with high sequence similarity, a “mismatch oligo” with one or two noncomplementary nucleotides (relative to the probe) was included on each membrane to test for probe specificity and cross-hybridization. There was clearly a signal of cross-hybridization to these mismatch oligos (which was expected given the permissive hybridization temperature of 48°C). Thus, the signal for the RNA samples likely reflects some additional hybridization for other isodecoders with very similar sequence at the probe region. There are multiple isodecoders that are similar in sequence to tRNA-ProTGG-112 and tRNA-SerCGA-3245 at the probe region, and the mismatch oligos were designed to be identical to these similar isodecoders. There are no similar tRNAs to GlyGCC-3370 and IleGAT-3373 (the mismatch oligos for these genes were therefore “synthetic” with no biological match to another gene). Therefore, the already low hybridization signal for tRNA-ProTGG-112 and tRNA-SerCGA-3245 is likely an overestimation of expression, further exacerbating the incongruence between the northern blot analysis and the massive tRNA-seq CPM values for tRNA-ProTGG-112.

### RT-induced misincorporations identify positions of base modifications in plant tRNAs

tRNA base modifications at the Watson-Crick face can interfere with the base pairing that is necessary for RT. Such modifications may not only stall or terminate reverse transcriptase activity but can also result in the misincorporation or deletion of nucleotides in the resulting cDNA. These misincorporations can be used to infer both the position and modification type, producing a modification map or “index” of a tRNA (Clark et al. 2016; Potapov et al. 2018; Vandivier et al. 2019). Reads were globally aligned to reference sequences to identify all nucleotide positions that differed from the reference gene **(supp. Table 9**). Even after treatment with AlkB, a signal of RT misincorporation was still present in almost one-third of the tRNAs in at least one position. We identified a position as confidently modified if ≥30% of the mapped reads varied from the reference sequence at that position. There was evidence of multiple tRNAs being modified at the same site, with the same position being modified in up to 17% of all reference tRNAs (**Figure 7)**. Given that AlkB may act on only a subset of modifications on certain bases (Clark et al. 2016), it was unsurprising that positions with a modified T were largely insensitive to AlkB treatment (**Table 1**). Similar to work with RT-based modification detection in human cell lines and yeast (Cozen et al. 2015; Zheng et al. 2015; Clark et al. 2016), we found that demethylation treatment with AlkB had a strong effect at only certain tRNA positions (e.g., 9, 26, and 58). The most frequent type of misincorporation differed by both position and reference base, but Gs were most likely to be deleted whereas the other three bases were most likely to be misread as a substitution **(Table 2).**

**Table 1.** Total proportion of reference nucleotides misread ≥30% of the time in *Arabidopsis thaliana* libraries

| Library | Reference |  |  |  |
| --- | --- | --- | --- | --- |
|  | A | C | G | T |
| AlkB+ (rep 1) | 0.0187 | 0.0028 | 0.0147 | 0.0149 |
| AlkB+ (rep 2) | 0.0168 | 0.0028 | 0.0113 | 0.0140 |
| AlkB+ (rep 3) | 0.0136 | 0.0011 | 0.0100 | 0.0112 |
| AlkB- (rep 1) | 0.0350 | 0.0038 | 0.0241 | 0.0100 |
| AlkB- (rep 2) | 0.0361 | 0.0032 | 0.0258 | 0.0089 |
| AlkB- (rep 3) | 0.0299 | 0.0036 | 0.0224 | 0.0098 |
| Untreated (rep 1) | 0.0347 | 0.0038 | 0.0237 | 0.0112 |
| Untreated (rep 2) | 0.0294 | 0.0032 | 0.0213 | 0.0086 |
| Untreated (rep 3) | 0.0339 | 0.0041 | 0.0213 | 0.0100 |

**Table 2.**
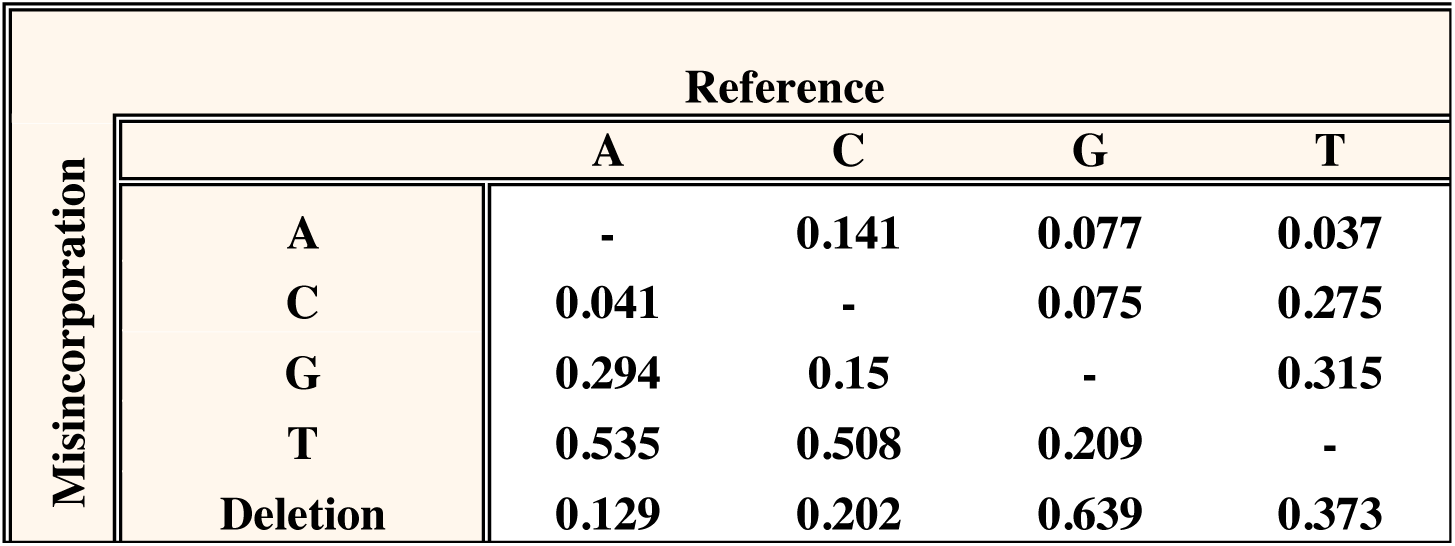
Total misincorporation frequency of tRNA reference bases across all nine *Arabidopsis thaliana* tRNA-seq libraries (see Table 1).

**Fig. 7.**
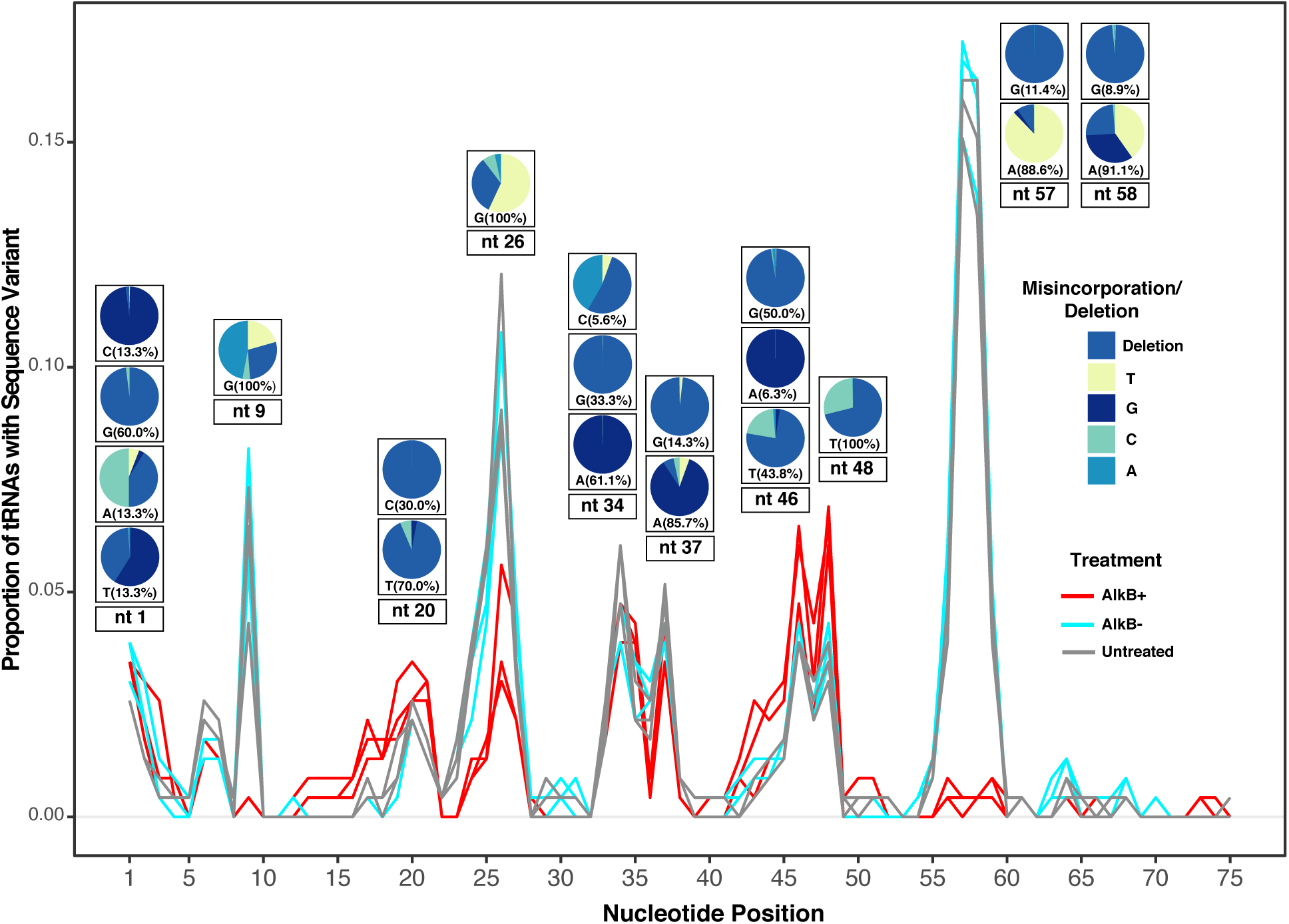
A tRNA modification index showing the proportion of all *A. thaliana* tRNA reference sequences with a misincorporation/deletion at each nucleotide position. A position was considered modified if ≥30% of the mapped reads differed from the reference sequence and the sequence was detected by more than five reads. Pie charts show the identity of the misincorporated base at some of the most frequently modified positions. A separate pie chart is provided for each observed reference base at that position, and the percentage indicates what proportion of modified tRNA sequences have that reference nucleotide. All three replicates of each treatment are shown and indicated by line color.

## DISCUSSION

### The promise and pitfalls of quantifying tRNA expression using tRNA-seq methods

In addition to the fundamental role of tRNAs as decoders of the genetic code, they are increasingly being recognized as integral players in a wide range of developmental, stress, tumorigenesis, biosynthetic, and amino acid delivery pathways (Hopper and Phizicky 2003; Banerjee et al. 2010; Phizicky and Hopper 2010; Chatrchyan et al. 2011; Raina and Ibba 2014; Kirchner and Ignatova 2015; Huang et al. 2018). As such, accurately detecting and quantifying tRNAs is key to gaining a more complete understanding of expression and regulation of numerous biological processes.

There have been substantial efforts in the past decade to make high-throughput sequencing methods more applicable to tRNAs. Nevertheless, we found that, even after treatment with AlkB and use of YAMAT adapters to specifically capture mature tRNAs, there still remained persistent RT-related sequencing bias, as evidenced by the extreme skew in tRNA-seq reads towards certain sequences (**Figures 3 and 4**). The observed dominance of multiple nuclear tRNA-Pro genes and a plastid tRNA-GlyGCC likely does not reflect biological reality, as there was no agreement between the tRNA-seq read abundance estimates and the intensity of hybridization in northern blot analysis, where tRNAs are probed directly without an intervening RT step. Moreover, the massive inferred expression level of plastid tRNA-GlyGCC is at odds with previous 2D-gel analysis of purified chloroplast tRNAs, which did not identify tRNA-GlyGCC as unusually abundant (Selden et al. 1983; Bergmann et al. 1984; Mubumbila et al. 1984). “Jackpot” tRNAs with extremely and presumably artefactually high read abundance have been reported in prior tRNA-seq results (Pang et al. 2014; Jacob et al. 2019), and the original YAMAT-seq paper reported a single tRNA-LysCTT gene having approximately 10-fold higher expression than the next most highly expressed gene (Shigematsu et al., 2017).

We found that four divergent angiosperm species showed the same general tRNA-seq profile dominated by nuclear tRNA-Pro and plastid tRNA-GlyGCC, suggesting an underlying cause of sequencing bias that is broadly shared across angiosperms. One possibility is that the overrepresented genes are less modified than other tRNAs. The plastid tRNA-GlyGCC and almost all of the nuclear tRNA-Pro isoacceptors had no RT misincorporations (based on our 30% threshold) after AlkB treatment. However, many other tRNAs lacked a strong signal for modification at any position (**supp. Table 9)** but did not show the same high abundance as nuclear tRNA-Pro and plastid tRNA-GlyGCC.

Modifications that predominantly result in the termination of RT (“RT falloff”) without other signatures of RT inhibition (i.e. base misincorporations and indels) are not detectable by the YAMAT-seq method that we employed because molecules are not sequenced unless RT proceeds all the way through the tRNA and captures the 5’ adapter. Thus, variation across different tRNAs in the presence of modifications that induce RT falloff could be responsible for observed RT bias. Additionally, the effect of certain modifications on RT behavior has been shown to be dependent on the nucleotide 3’ of modifications in the template RNA (Hauenschild et al. 2015), suggesting that even if a modification is detected and identified, it may have a different effect on expression analysis depending on the tRNA sequence. Finally, RT of tRNAs is also affected by primary and secondary structure (Motorin et al. 2007). Thus, there may be otherwise ubiquitous secondary structure characteristics that are not present in plastid tRNA-GlyGCC or multiple nuclear tRNA-Pro isoacceptors, making these tRNAs more amenable to RT. These represent important areas of investigation to further understand and alleviate sources of bias in tRNA-seq methods.

The biases that we have identified make it clear that more work must be done in developing a tRNA sequencing method that can accurately quantify *absolute* levels of expression. Nevertheless, the combination of YAMAT-seq and AlkB treatment has a number of advantages and great promise for analyzing changes in *relative* tRNA expression across treatments, tissues or subcellular fractions. In particular, we were able to generate a high proportion of full-length tRNA reads which can be confidently mapped to a single reference gene. Mapping of tRNA-seq reads to loci can be problematic because of the large number of similar but non-identical tRNA gene sequences. Other tRNA sequencing methods that hydrolyze tRNA [Hydro-tRNA-seq] (Gogakos et al. 2017), or utilize a template-switching reverse transcriptase [DM-TGIRT-seq] (Zheng et al. 2015) produce a large proportion of tRNA fragments that can ambiguously map to multiple genes. As there is increasing attention on the regulation and expression of specific tRNA genes (Hummel et al. 2019; Torres et al. 2019), the combination of tRNA-seq methods that we employed could be effectively used for differential expression analysis when trying to tease apart the transcriptional activity and turnover of individual genes within large gene families. Looking forward, technologies that directly analyze RNA molecules without the intermediate step of RT, such as nanopore sequencing (Jain et al. 2016), offer promise to accurately quantify tRNA expression without the confounding effects of RT bias. However, nanopore technologies still require substantial development to achieve sufficient sequencing accuracy and differentiation of tRNA species, especially in the context of the extensive base modifications present in tRNAs (Smith et al. 2015). In the interim, our combined approach may represent one of the more effective means to quantify relative tRNA expression changes at a single-gene level.

### The landscape of base modifications in plant tRNAs

Although base modifications likely contribute to biased quantification of tRNA expression, there remains a large benefit of utilizing a RT-based tRNA-seq methods because RT misincorporation behavior provides insight into the location and identity of base modifications. In addition to modifications that terminate cDNA synthesis, RTs can read through methylations at the Watson-Crick face with varying efficiencies, resulting in misincorporations and indels in the cDNA (Tserovski et al. 2016; Pan 2018; Potapov et al. 2018). Here, we took advantage of RT-based expression analysis to present one of the most extensive modification landscapes of angiosperm tRNAs to date. The modification peaks produced from sequencing complete *A. thaliana* tRNAs (**Figure 7**) provide an informative comparison to the annotated modification indexes previously generated for human tRNAs (Clark et al. 2016). Given the widely conserved presence of some tRNA modifications known to inhibit RT (Jackman and Alfonzo 2013), it was unsurprising that the modification indexes from plants and humans shared many similarities, including a high rate of modification at nucleotide positions 9, 20, 26, 34, 37, and 58 (**Figure 8**). We reported modifications based on their actual position in tRNA genes. Thus, in some cases, the same modification at functionally analogous sites in different tRNAs can be represented by peaks spanning two or three nt positions due to differences in tRNA length.

**Fig. 8.**
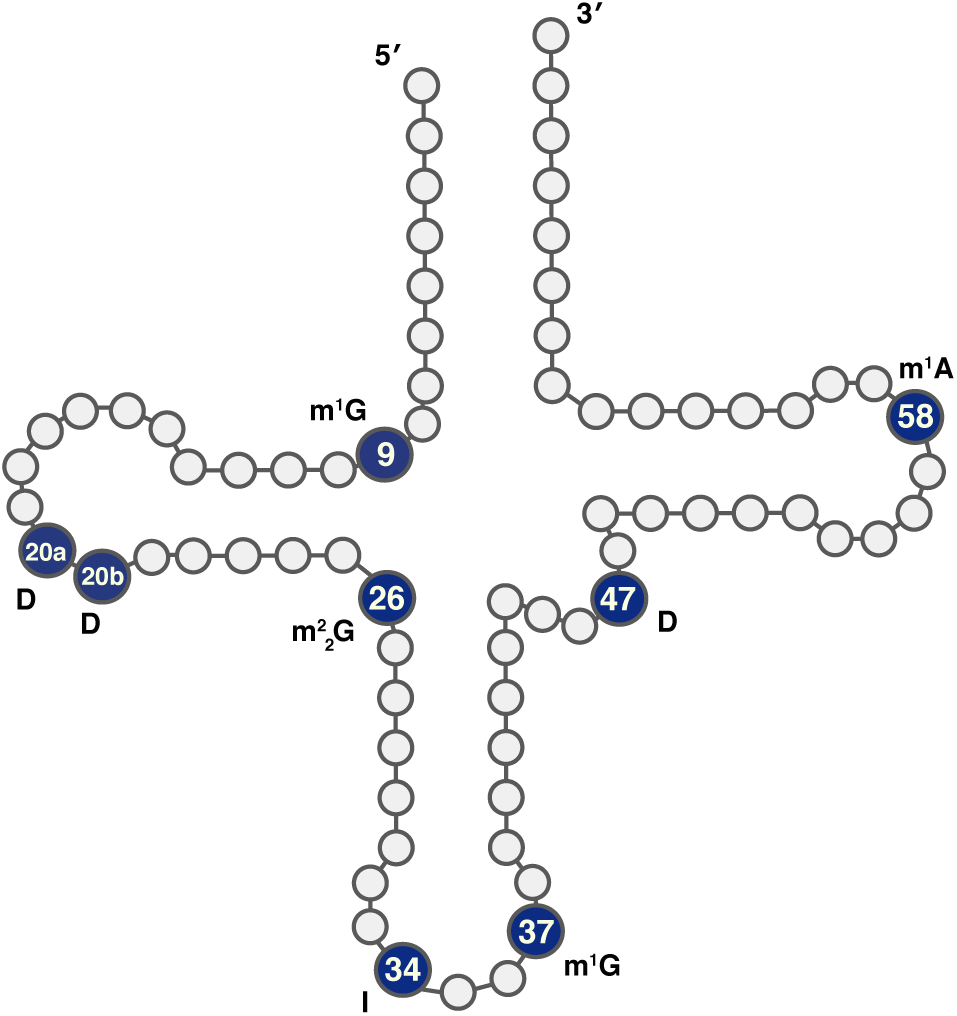
tRNA positions labeled with possible modifications. Positions with strong misincorporation signals are indicated with a known modification at that position in other species. D: dihydrouridine; I: inosine; m^1^A: N1-methyladenosine; m1G: N1-methylguanosine; m^2^_2_G: N2,N2-dimethylguanosine

Previous work (Zheng et al. 2015) has confirmed position 9 as frequently having an m^1^G modification and position 58 having an m^1^A modification. These modifications are known targets of AlkB demethylation, and we found that AlkB treatment resulted in almost complete reduction of the modification index peaks at positions 9 and 58 (some of this signal appears at position 57 in our modification index because of differences in tRNA length) (**Figure 7**), adding support for the existence of these same modifications in plants. Two consecutive dihydrouridines are commonly found in the D loop around position 20 in eukaryotes (Hendrickson 2001), a modification motif that may also be common in plant tRNAs, as we found a modification peak around positions 19-21. Position 26 is often modified to N2,N2-dimethylguanosine (m^2^_2_G), which also appears to be mildly sensitive to AlkB demethylation treatment (Clark et al. 2016). We found a modification peak at position 26 and a small reduction in RT misincorporation rate after AlkB treatment in *A. thaliana*. Adenosine-to-inosine modifications are frequently found at position 34 in eukaryotic tRNAs (Torres et al. 2015). RT of inosine is expected to cause A-to-G changes because inosine base pairs with C instead of T (Motorin et al. 2007). Indeed, we found almost 100% misincorporation of G when the reference was A at position 34. It is important to note that some tRNA modifications may not be present in all copies (Arimbasseri et al. 2015) or may not always cause RT misincorporation (Potapov et al. 2018). Thus, our modification index with a cutoff of ≥30% misincorporation is focusing on highly modified positions in the corresponding tRNAs (Clark et al. 2016).

Our analysis also identified clear differences in the modification index between plants vs. humans. In particular, we found a strong misincorporation peak at positions 46-48 for T reference nucleotides that was not observed in humans (**Figure 8**) (Clark et al. 2016). The misread Ts at position 46-48, could correspond to dihydrouridines, which are commonly found in the variable loop (position 47 in standard tRNA nomenclature) of eukaryotic and plastid tRNAs (Xing et al. 2004; Lorenz et al. 2017). Taken together, over 14% of all *A. thaliana* reference tRNAs appear to have a modified T at the position, including all cytosolic and plastid tRNA-Ile isoacceptors, as well as all cytosolic tRNA-ThrAGT, tRNA-AsnGTT, and tRNA-TyrGTA isodecoders. This trend of frequent modification at position 47 was conserved in all angiosperm species tested (**supp. Figure 3, supp. Table 10**). It is interesting to note that the misincorporation profile (deletions and T-to-C misincorporations) for dihydrouridines at positions 19-21 is similar to the misincorporation profiles that we observe for positions 46-48. Many *A. thaliana* tRNAs have a G in their reference sequence at position 46 and frequently exhibited a deletion at this site. This pattern likely reflects a modification to m^7^G, which is predicted to be common in cytosolic Viridiplantae tRNAs at that position (Machnicka et al., 2014). Adding to this prediction is the complete lack of an AlkB effect on the modification peak at position 46, as AlkB is not expected to remove m^7^G modifications. Our analysis demonstrates how tRNA-seq methods can be used to elucidate similarities and differences in global tRNA modification patterns between divergent taxonomic groups. These approaches could be extended to assess whether different tissues or even subcellular compartments depend more heavily on certain tRNA modifications for functionality.

In addition to generating a general modification index across the entire tRNA population, this tRNA-seq method can be used to identify the modification profiles of individual tRNAs or specific groups of tRNAs. In other eukaryotes, organellar tRNAs have been found to be less modified than their cytosolic counterparts (Machnicka et al. 2014; Salinas-Giege et al. 2015). In agreement with this, we detected fewer modifications on average in *A. thaliana* organellar tRNAs. Cytosolic tRNAs with a misincorporation signal had an average of 3.0 modifications in AlkB-libraries, whereas organelle tRNAs with misincorporations had an average of 2.1 modifications. Likely owing to greater rates of modification in cytosolic tRNAs, AlkB treatment had a substantially larger effect on improving the detection of cytosolic tRNAs than on organellar tRNAs (**Figure 2**). However, some organellar tRNAs exhibited the greatest extent of modification as detected by RT-based sequence changes. The plastid tRNA-IleCAT had the most positions exhibiting a misincorporation signature (16) even after AlkB treatment, and mitochondrial tRNA-fMetCAT was a close second with 14 such positions prior to demethylation.

It is sometimes assumed that tRNAs with the same anticodon but different body sequences will have similar modification patterns (Torres et al. 2019). Nevertheless, we found examples of large differences in modification patterns among isodecoders. For example, the large nuclear tRNA-GlyGCC isodecoder family has a range of one to six inferred modifications depending on the reference sequence and show a range of sensitivity to AlkB treatment. That some tRNAs, even those with the same anticodon, have such radically different modification patterns represents an interesting and largely unexplored facet of tRNA biology, and offers support to the finding that isodecoders may differ in function (Geslain and Pan 2010).

We found that AlkB treatment sometimes created novel RT misincorporations instead of restoring the correct nucleotide. For example, the C in the anticodon of the plastid tRNA-IleCAT is known to be modified to lysidine, which has base-paring behavior like that of U (Kashdan and Dudock 1982; Alkatib et al. 2012), and we found this modification to be consistently misread as a deletion in AlkB+ libraries, whereas it was detected as a T, or even rarely as a C, in untreated libraries. Additionally, an increase in abundance of some tRNA fragments was found in AlkB+ libraries (**supp. Table 11**) and spatially associated with modified bases. For example, the “U-turn” fragments generated from tRNA-ThrCGT-3316 invert at what appears to be a modified G25, and the abundance of these fragments increased with AlkB treatment.

Historically, comprehensive mapping of base modifications has been laborious and depended on tRNA purification followed by digestion and mass spectrometry (Su et al. 2014). Quickly and accurately identifying tRNA modifications through RT-based methods makes it possible to test hypotheses regarding the presence and function of modifications at phylogenetic, developmental, and subcellular levels. tRNAs are by far the most post-transcriptionally modified gene class (Boccaletto et al. 2018). Research is just now starting to tease apart the many roles these modifications play (Bjork et al. 2001; Phizicky and Alfonzo 2010; Pereira et al. 2018; de Crecy-Lagard et al. 2019). Not all modifications are detectable through sequencing, but it has been estimated that RT-based methods can identify approximately 25% of tRNA modifications in humans (Clark et al. 2016), and the modifications that can be detected by sequencing are known to have critical roles in tRNA folding, rigidity and stability (Pan 2018). Given the diversity of tRNA modifications, the full scope of RT behavior and the effect of AlkB treatment on tRNA base modifications has yet to be fully explored, but work is currently being done to further characterize RT behavior when encountering specific modifications (Ebhardt et al. 2009; Potapov et al. 2018). The misincorporation signatures resulting from the RT of modified bases are also being used in machine-learning algorithms to identify new modifications (Vandivier et al. 2019). Moving beyond RT-based methods to direct sequencing of tRNAs and their modifications is another exciting development on the horizon. However, major advances must still be made in techniques such as nanopore sequencing because current technologies cannot be used to reliably sequence tRNA molecules in biological total RNA samples because they are likely unable to discriminate between more than two tRNAs at a time (Henley et al. 2016; Poodari 2019). Additionally, efforts to characterize the resistance properties of alternative tRNA base modifications in nanopores are underway (Onanuga et al. 2017) but still in their infancy. Thus, along with the promise of additional technologies on the horizon, current RT-based methods offer easily accessible and exciting tools to study base modification profiles of full-length tRNA sequences and how they vary across taxa, treatments, tissues, and subcellular locations.

## METHODS

### Plant material and growth conditions

For tRNA-seq material, *Arabidopsis thaliana* Columbia ecotype (Col-0) was grown in Pro-Mix BX General Purpose soil supplemented with vermiculite and perlite. Plants were germinated and kept in growth chambers at Colorado State University 23°C with a 16-hr/8-hr light/dark cycle (light intensity of 100 μE×m^-2^×s^-1^). *Solanum tuberosum* var. Gladstone (PI: 182477), *Oryza sativa* var. Ai-Chiao-Hong (PI:584576) and *Medicago truncatula* (PI:660383) were acquired from the U.S. National Plant Germplasm System (https://www.ars-grin.gov/npgs/). Tissue culture plantlets of *S. tuberosum* were transferred to Pro-Mix BX General Purpose soil supplemented with vermiculite and perlite and kept in growth chambers with the same settings as above. Seeds of *O. sativa*, and *M. truncatula* were germinated in SC7 Cone-tainers (Stuewe and Sons) on a mist bench under supplemental lighting (16-hr/8-hr light/dark cycle) in the Colorado State University greenhouse, then moved to a growth chamber with the same settings as above after 4 weeks. For northern blot analysis, *A. thaliana* Col-0 was grown in a 12-hr/12-hr light/dark cycle in growth chambers at the Institute of Molecular Biology of Plants.

### RNA extraction

For our primary tRNA-seq experiment and ddPCR analysis, RNA from *A. thaliana* and *O. sativa* was extracted from 7-week-old leaf tissue. RNA from *M. truncatula* was extracted from 3-week-old leaf tissue, and RNA from *S. tuberosum* plantlets was extracted from leaf tissue 3 weeks after transfer into soil. Extractions were performed using a modified version of the Jordon-Thaden et al. (2015) protocol. In brief, tissue was frozen in liquid nitrogen and pulverized with a mortar and pestle and then vortexed and centrifuged with 900 μl of hexadecyltrimethylammonium bromide (CTAB) lysis buffer supplemented with 1% polyvinylpyrrolidone (PVP) and 0.2% β-mercaptoethanol (BME). Samples were then centrifuged, and the aqueous solution was removed. 800 μL of a chloroform:isoamyl alcohol (24:1) solution was added, mixed by inverting and centrifuged. The aqueous phase was removed, and 900 μl of TRIzol was added. The solution was then mixed by inversion and centrifuged. The aqueous phase was removed, 200 μl of chloroform was added, and the solution was then mixed and centrifuged again. The aqueous phase was removed followed by isopropanol RNA precipitation, and cleaned pellets were resuspended in dH_2_O. RNA was checked for integrity with a TapeStation 2200 and purity on a Nanodrop 2000. Later RNA extractions from *A. thaliana* for the AlkB D135S tRNA-seq experiment (8-week-old tissue) and northern blot analysis (4-week-old tissue) were performed with a simplified protocol. Leaf tissue was frozen in liquid nitrogen and pulverized with a mortar and pestle prior to the addition of TRIzol but otherwise followed the TRIzol manufacturer’s RNA extraction protocol. Three biological replicates (different plants) were used for the *A. thaliana* experiments, and a single sample was used for each of the three other angiosperms.

### AlkB purification

Plasmids containing cloned wild type AlkB protein (pET24a-AlkB deltaN11 [plasmid #73622]) and D135S mutant protein (pET30a-AlkB-D135S [plasmid #79051]) were obtained from Addgene (http://www.addgene.org/). Protein was expressed and purified at CSU Biochemistry and Molecular Biology Protein Expression and Purification Facility. Cells were grown at 37°C to an OD_600_ of 0.6, at which time, 1 mM of isopropyl-β-D-thiogalactoside was added, and temperature was lowered to 30°C. Cells were harvested after 3 hr by centrifugation and resuspended in 10 mM Tris (pH 7.3), 2 mM CaCl_2_, 300 mM NaCl, 10 mM MgCl_2_, 5% glycerol, and 1 mM BME. Resuspension was homogenized by sonication, and lysate was recovered by centrifugation. The supernatant was loaded onto HisTrap HP 5 ml columns (GE Healthcare) and was washed and eluted by a linear gradient of 0-500 mM imidazole. The fractions containing AlkB were pooled and concentrated with ultrafiltration using Amicon Ultra-15 MWCO 10kDa (Millipore). The concentrated sample was then loaded onto HiLoad 16/60 Superdex 200 prep grade size exclusion column (GE Healthcare) in 20 mM Tris pH 8.0, 200 mM NaCl, 2 mM DTT, and 10% glycerol. The fractions containing AlkB were pooled and concentrated, aliquoted, flash-frozen in the presence of 20% glycerol and stored at −70°C.

### Demethylation reaction

AlkB reactions were performed using a modified version of existing protocols (Cozen et al. 2015; Zheng et al. 2015; Chen et al. 2016). Demethylation was performed by treating 10 μg of total cellular RNA with 400 pmols of AlkB in a reaction volume of 80 μl containing: 70 μM Ammonium Iron(II) Sulfate Hexahydrate, 0.93 mM α-Ketoglutaric acid disodium salt dihydrate, 1.86 mM ascorbic acid, and 46.5 mM HEPES (pH 8.0), incubated at 37°C for 60 min. The reaction was quenched by adding 4 μl of 100 mM EDTA followed by a phenol-chloroform RNA extraction, ethanol precipitation with the addition of 0.08 μg of RNase-free glycogen, and resuspension in water. RNA integrity was checked on a TapeStation 2200. The same procedure was followed in parallel for AlkB-control libraries except that the AlkB enzyme in the reaction volume was replaced with dH_2_O.

### Illumina tRNA-seq library construction

All adapter and primer sequences used in library construction can be found in supplementary Table 12. In order to remove amino acids from the mature tRNAs (deacylation), demethylated or control RNA was incubated in 20 mM Tris HCl (pH 9.0) at 37°C for 40 min. Following deacylation, adapter ligation was performed using a modified protocol from (Shigematsu et al. 2017). A 9 μl reaction volume containing 1 μg of deacylated RNA and 1 pmol of each Y-5’ adapter (4 pmols total) and 4 pmols of the Y-3’ adapter was incubated at 90°C for 2 min. 1 μl of an annealing buffer containing 500 mM Tris-HCl (pH 8.0) and 100 mM MgCl_2_ was added to the reaction mixture and incubated for 15 min at 37°C. Ligation was performed by adding 1 unit of T4 RNA Ligase 2 enzyme (New England Biolabs) in 10 μl of 1X reaction buffer and incubating the reaction at 37°C for 60 min, followed by overnight incubation at 4°C. We found that adapter ligation with the input of 80 pmols of adapters called for in the original protocol resulted in an excess of adapter dimers and other adapter-related products after PCR and that using 1/10^th^ of that adapter concentration maximized yield in the expected size range for ligated tRNA products, while minimizing adapter-related products.

RT of ligated RNA was performed using SuperScript IV (Invitrogen) according to the manufacturer’s protocol. Briefly, 1 μl of 2 μM RT primer, and 1 μl of 10 mM dNTP mix was added to 11 μl of the deacylated RNA from each sample. The mixture was briefly vortexed, centrifuged and incubated at 65°C for 5 min. Then, 4 μl of 5X SSIV buffer, 1 μl 100 mM DTT, 1 μl RNaseOUT, and 1 μl of SuperScriptIV were added to each reaction. The mixture was then incubated for 10 min at 55°C for RT and inactivated by incubating at 80°C for 10 min.

The resulting cDNA was then amplified by polymerase chain reaction (PCR) in a 50 μl reaction containing 7 μl of the RT reaction, 25 μl of the NEBNext 2X PCR Master Mix, 1 μl of the PCR forward primer, 1 μl of the PCR reverse primer, and 15.5 μl dH_2_O. Ten cycles of PCR were performed on a Bio-Rad C1000 Touch thermal cycler with an initial 1 min incubation at 98°C and 10 cycles of 30 s at 98°C, 30 s at 60°C and 30 s at 72°C, followed by 5 min at 72°C.

Size selection of the resulting PCR products was done on a BluePippin (Sage Science) with Q3 3% agarose gel cassettes following the manufacturer’s protocol. The size selection parameters were set to a range of 180-231 bp, with a target of 206 bp. Size-selected products were then cleaned using solid phase reversable immobilization beads and resuspended in 10 mM Tris (pH 8.0).

### Sequencing and read processing

The nine original *A. thaliana* tRNA-seq libraries (three AlkB+, three AlkB-, and three entirely untreated) were single-indexed and sequenced on an Illumina MiSeq with single-end, 150 bp reads. Libraries from *M. truncatula, O. sativa, S. tuberosum*, and the *A. thaliana* D135S AlkB mutant and wild type AlkB libraries were dual-indexed and sequenced on an Illumina NovaSeq 6000 with paired-end, 150-base pair reads. Sequencing reads are available via the NCBI Sequence Read Archive under BioProject PRJNA562543. Adapters were trimmed using Cutadapt version 1.16 (Martin 2011) with a quality-cutoff parameter of 10 for the 3’ end of each read. A minimum length filter of 5 bp was applied to reads from the MiSeq sequencing platform. Read length filters of a minimum of 50 bp and a maximum of 95 bp were applied to reads produced from the NovaSeq 6000, as a much larger percentage of the reads from those libraries were <20 bp after adapter trimming. For paired-end data, BBMerge from the BBTools software package was used to merge R1 and R2 read pairs into a consensus sequence (Bushnell et al. 2017). Identical reads were summed and collapsed into read families using the FASTQ/A Collapser tool from the FASTX-Toolkit version 0.0.13 (http://hannonlab.cshl.edu/fastx_toolkit/index.html).

### Sequence alignments and mapping

Mapping for all processed and collapsed reads was performed with a custom Perl script using reference tRNA databases from the PlantRNA website (http://seve.ibmp.unistra.fr/plantrna/). Each collapsed read family was BLASTed (blastn, 1e-6) against a complete set of (non-identical) nuclear, mitochondrial and plastid reference tRNA sequences from the corresponding species, and the best match was assigned to each read family. The read count of all assigned read families was then summed for each reference sequence. When a read family was an equally good match to multiple references, the read count was divided by the total number of tied references. Multiple BLAST statistics were recorded for each read family/reference hit pair including e-value, hit score, hit length, and percent identity.

Because we were mainly interested in mature tRNA sequences, only reads that had 80% or greater hit coverage to a reference sequence were used in downstream analysis. Complete datasets including reads that fell below the 80% hit coverage threshold were used to determine read length distributions and the proportion of reads that did not BLAST to a tRNA reference sequence.

### Generation of modification index

In order to capture all possible misincorporations and indels including those that could have occurred at the end of the reads, a custom Perl script and the alignment software MAFFT version 7.407 (Katoh et al. 2017) was used to produce full-length alignments of reads to their top BLAST reference tRNA hit and identify all base substitutions and indels in the resulting alignment. Only reads that BLASTed to their reference sequence with an e-value of 1e-10 or less and had 90% hit coverage to the reference were retained for modification index calling. Only positions that were covered by greater than five reads were used for modification index calling, and a position was considered confidently modified if ≥30% of the mapped reads at that position differed from the reference nucleotide by either a substation or deletion. Scripts used for read processing, mapping, and sequence analysis are available via GitHub (https://github.com/warrenjessica/YAMAT-scripts).

### ddPCR

To generate cDNA for ddPCR quantification, 1 μg of RNA from each of the three original AlkB+ *A. thaliana* samples was treated with DNase I (Invitrogen) according to the manufacturer’s protocol. For each sample, 146 ng of DNase-treated RNA was then subjected to RT using iScript Reverse Transcription Supermix (BioRad) according to manufacturer’s protocol in a 20-μl reaction volume. cDNA was then diluted in a four step concentration series to 0.001ng/μl, 0.01 ng/μl, 0.1 ng/μl, and 1 ng/μl. ddPCR was performed with each of the four primer pairs (**supp. Table 6**) using this concentration series of template cDNA to determine ideal concentration. All ddPCR amplifications were set up in 20-μL volumes with Bio-Rad QX200 ddPCR EvaGreen Supermix and a 2 μM concentration of each primer before mixing into an oil emulsion with a Bio-Rad QX200 Droplet Generator. The final template input amounts were 0.1 ng for amplification of tRNA-GlyGCC-3370 and tRNA-CysGCA-3354, 0.001 ng for amplification of tRNA-ProTGG-112, and 1 ng for amplification of tRNA-GlyCCC-93. Amplification was performed on a Bio-Rad C1000 Touch Thermal Cycler with an initial 5 min incubation at 95°C and 40 cycles of 30 s at 95°C and 1 min at 60°C, followed by signal stabilization via 5 min at 4°C and 5 min at 95°C. The resulting droplets were read on a Bio-Rad QX200 Droplet Reader. Copy numbers for each PCR target were calculated based on a Poisson distribution using the Bio-Rad QuantaSoft package.

### Northern blots

RNAs and synthesized oligonucleotides (**supp. Table 7**) were separated on 15% (w/v) polyacrylamide gels. Gels were then electrotransfered onto Hybond-N^+^ nylon membranes (Amersham) for 90 min at 300mA/250V and UV-crosslinked. All membranes were hybridized with ^32^P-radiolabeled oligonucleotide probes (**supp. Table 7**) at 48°C in a 6X saline-sodium citrate (SSC) buffer with 0.5% sodium dodecyl sulphate (SDS) solution for 14 hr. Hybridization was followed by two washes (10 min) with 2X SSC buffer at 48°C buffer followed by one wash (30 min) at 48°C in 2X SSC with 0.1% SDS. Membranes were imaged on a Typhoon FLA 9500 biomolecular imager (GE Healthcare Life Sciences) after 16 hr of exposure on film.

Northern blots were analyzed using ImageJ, version Java 1.8.0_172 (64-bit) (https://imagej.nih.gov/ij/) to quantify signal intensity for each band. Signal peaks were quantified by first drawing a straight (0.00 degree) line beginning at the farthest left point of contact of the signal to the opposite side of the window. To define only the major signal peak of interest, two lines were drawn perpendicular to the horizontal. The position of these lines was determined by eye based on where the peak’s curves began to smooth. The coordinates of these vertical boundaries were used for all subsequent plots of the same membrane. The area was then calculated between the horizontal line and the peak signal bounded by the two vertical lines. Using the known concentration of the loaded oligos and the corresponding signal area, the estimated pmols of each tRNA target was calculated by fitting a standard curve to the log_10_ values of signal intensity and oligo concentration (**supp. Table 7**).

### Statistics and figure generation

Differential expression analysis between the AlkB treated and untreated libraries in the original analysis as well as the wild type and D135S mutant AlkB libraries was done with the R package EdgeR version 3.24.3 (Robinson et al. 2010), using a dataset with only reads that had 100% hit coverage to a reference sequence. A linear regression was performed in R with the lm function to test the relationship between tRNA-seq CPM values and ddPCR copy-number estimates per ng of input RNA.

## ACKNOWLEDGMENTS

We thank Deyu Li for providing valuable guidance in optimizing the AlkB reaction and all of the research personnel at the Institute of Molecular Biology of Plants for graciously hosting J.M.W. We would also like to thank Dr. Carol Wilusz and Adam Heck for test RNA samples and valuable discussions, and Dr. John Hunt and Dr. Victor Wong at Columbia University for sending samples of AlkB for initial testing. This work was supported by a Chateaubriand Fellowship from the Embassy of France in the United States, an NSF GAUSSI graduate research fellowship (DGE-1450032), NSF award IOS-1829176, and start-up funds from Colorado State University.

---

**Supp. Fig. 1.**
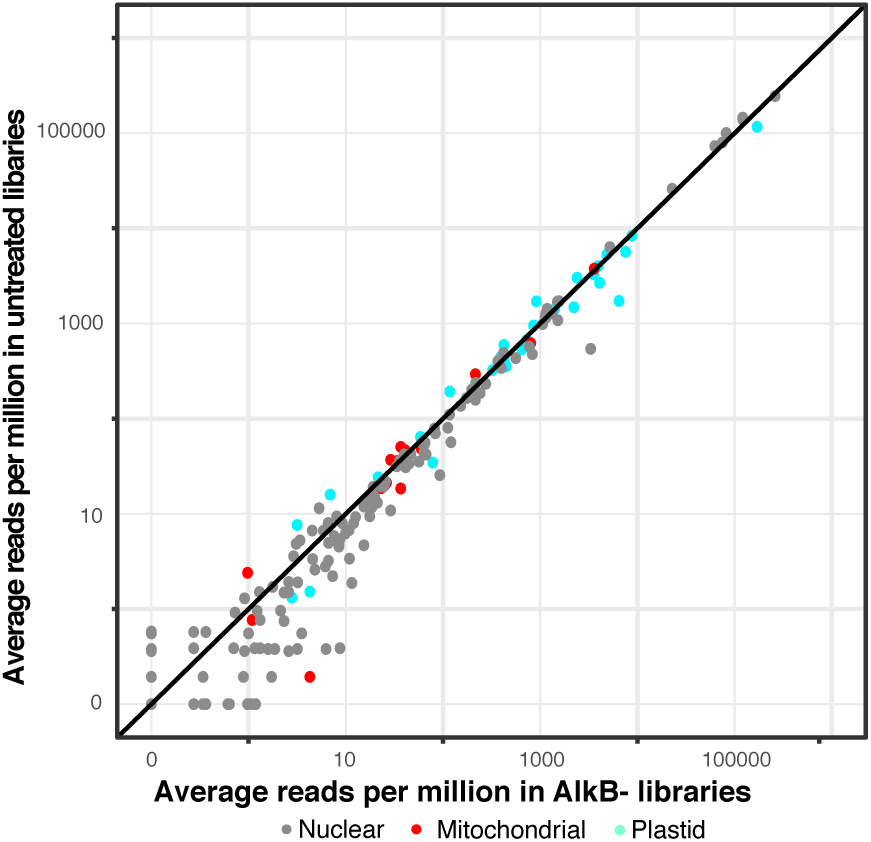
Average read counts per million across three biological replicates are shown for each unique *A. thaliana* tRNA reference sequence for untreated libraries and AlkB-(buffer only) libraries.

**Supp. Fig. 2.**
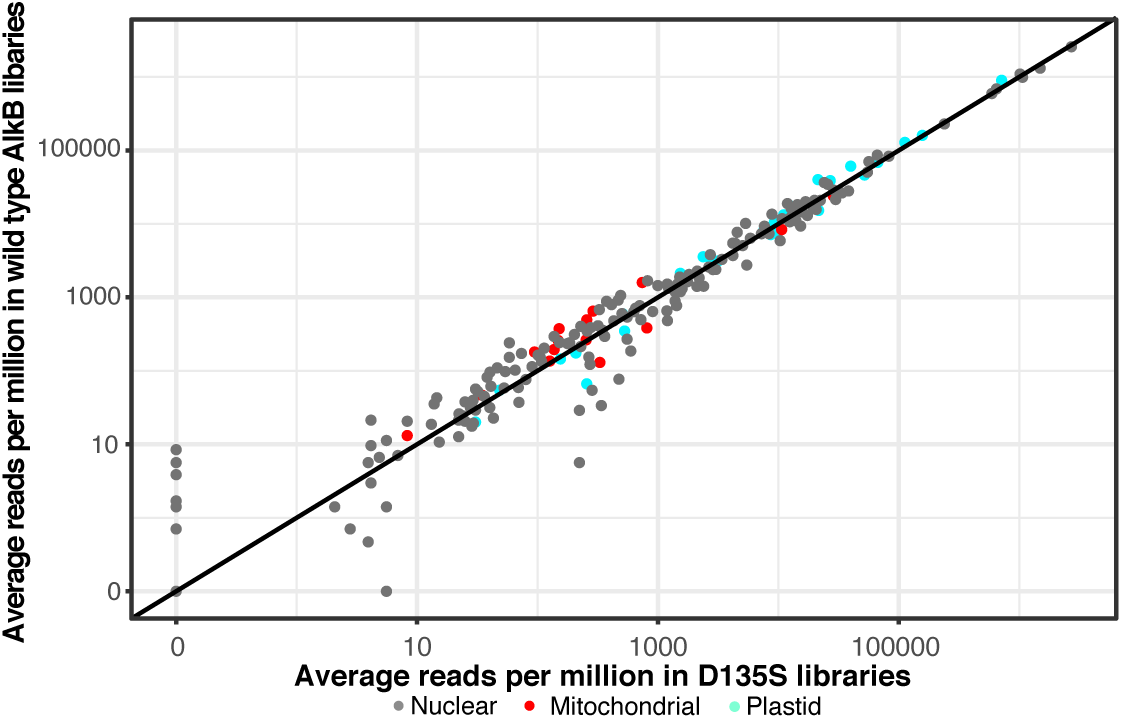
Average read counts per million across three biological replicates are shown for each unique *A. thaliana* tRNA reference sequence for wild type AlkB and D135S AlkB libraries.

**Supp. Fig. 3.**
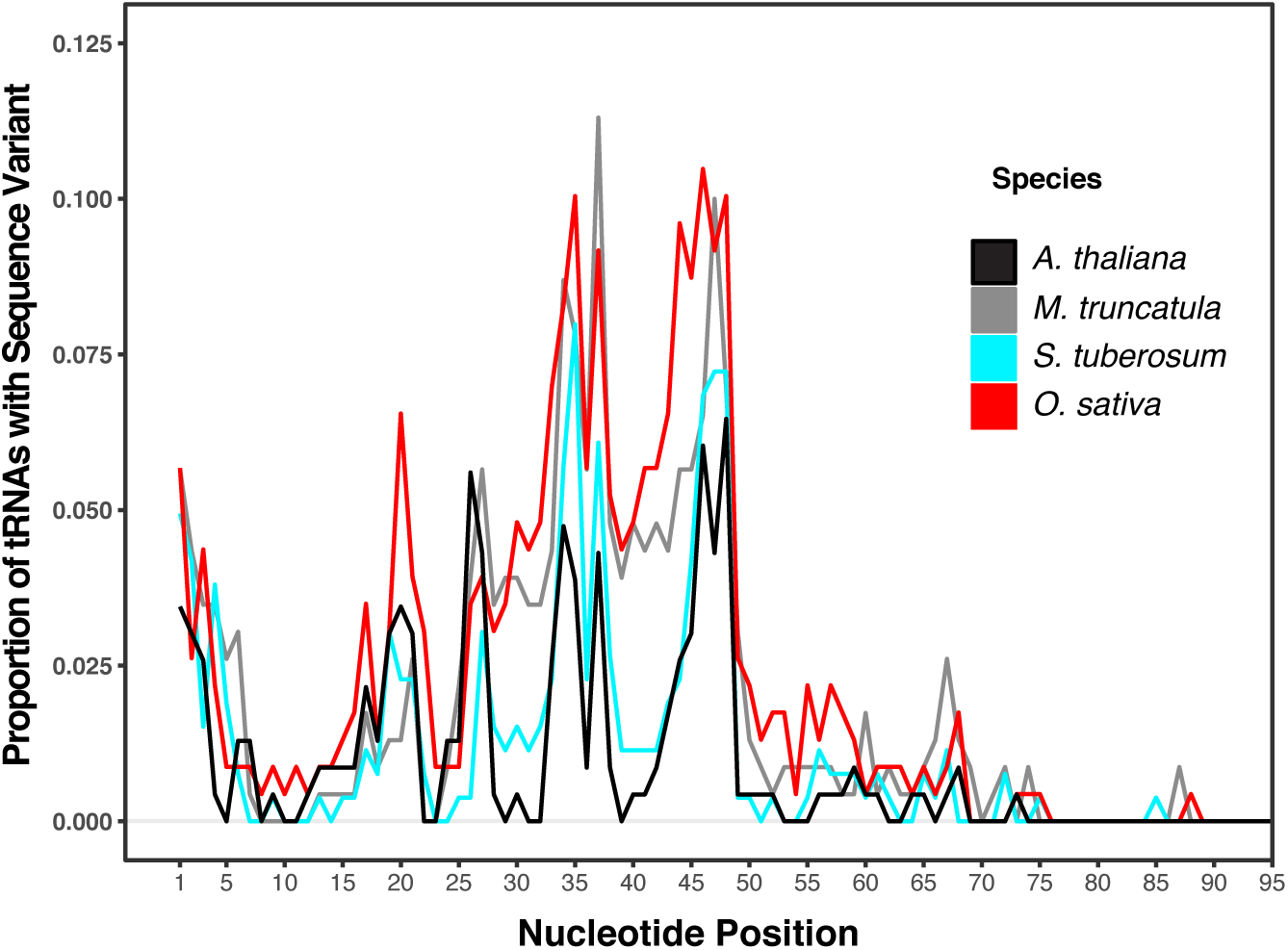
A tRNA modification index showing the proportion of *A. thaliana, M. truncatula, S. tuberosum, and O. sativa* tRNA reference sequences with a misincorporation/deletion at each nucleotide position. A position was considered modified if ≥30% of the mapped reads differed from the reference sequence and the sequence was detected by more than five reads. The modification index for *A. thaliana* was produced from the AlkB+ (rep 1) library.

